# Software Application Profile: mrrobust - a tool for performing two-sample summary Mendelian randomization analyses

**DOI:** 10.1101/142125

**Authors:** Wesley Spiller, Neil M. Davies, Tom M. Palmer

## Abstract

**Motivation:** In recent years Mendelian randomization analysis using summary data from genome-wide association studies has become a popular approach for investigating causal relationships in epidemiology. The mrrobust Stata package implements several of the recently developed methods.

**Implementation:** mrrobust is freely available as a Stata package.

**General Features:** The package includes inverse variance weighted estimation, as well as a range of median and MR-Egger estimation methods. Using mrrobust, plots can be constructed visualising each estimate either individually or simultaneously. The package also provides statistics such as 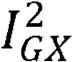 which are useful in assessing attenuation bias in causal estimates.

**Availability:** The software is freely available from GitHub [https://raw.github.com/remlapmot/mrrobust/master/].

## Introduction

Mendelian randomization^1^ has developed into a popular approach to examining causal relationships in epidemiology^2, 3^. By employing genetic variants as instrumental variables (IVs) it is possible to limit bias from confounding, provided variants satisfy the assumptions of IV analysis^1, 4^ For a genetic variant to serve as a suitable instrument, three assumptions must hold, 1) it must be associated with the exposure of interest, 2) there must be no confounders of the instrument and outcome, and 3) the instrument must not affect the outcome except via the exposure of interest.^5^.

Candidate variants are usually identified through large genome-wide association studies (GWASs) ^6^. However, IV analyses using a single variant rarely have sufficient power to test hypotheses of interest^6, 7^ One approach to increase the statistical power of Mendelian randomization studies is to use multiple genetic variants as instruments within a two-sample summary framework^8, 9^ Two-sample Mendelian randomisation estimates the effect of the exposure using instrument-exposure and instrument-outcome associations from different samples, often through methods originally developed for meta-analysis^8, 9^ This is particularly useful, as MR estimators such as MR Egger and median based regression are robust to certain forms of violation of the third instrumental variable assumption^8, 10, 11^. Violations of this assumption can occur through directional pleiotropy-where a genetic variant affects the study outcome through pathways that are not mediated via the exposure. Such developments have contributed to the increasing popularity of two-sample summary MR^5^.

This paper introduces the mrrobust Stata package as a tool for performing two-sample summary MR analyses. The mrrobust package is a tool to help researchers implement two-sample MR analyses, and can be viewed as the Stata counterpart to toolkits such as the MendelianRandomization R package^12^. Before continuing, we briefly outline the three primary estimation methods included in the mrrobust package, using the notation of Bowden et al^10^,^13^.

### Inverse variance weighting (IVW)

To perform IVW a weighted average 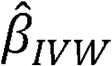 is calculated using the set of ratio estimates 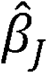 for each individual variant *J*= 1,2,…,*j* ^9^ Let 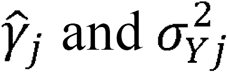 denote the instrument-outcome association and variance respectively for the *j^th^* variant. The IVW estimate is then defined as:

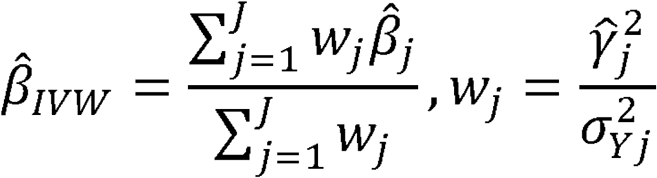

This corresponds to the estimate one would obtain from a weighted linear regression of the set of instrument-outcome associations upon the set of instrument-exposure associations, constraining the intercept at the origin^9^. One drawback of the IVW approach is that causal effect estimates can be biased in cases where one or more variants exhibit directional pleiotropy^9^.

### MR-Egger regression

MR-Egger regression is valid under weaker assumptions than IVW, as it can provide unbiased causal effect estimates even if the variants have pleiotropic effects.

In this case, the set of instrument-outcome associations is regressed upon the set of instrument-exposure associations, weighting the regression using precision of the instrument-outcome associations as in the IVW case^8^. However, MR-Egger does not constrain the intercept at the origin, and the intercept represents an estimate of the average directional pleiotropic effect across the set of variants. The slope of the model provides an unbiased estimate of the causal effect ^8 10^. If there is little evidence of systematic differences between the IVW and MR-Egger, then the IVW should be preferred. The IVW is more efficient, but potentially less robust, and in such cases the IVW estimate is often most appropriate estimate to adopt due to the greater precision of IVW estimates in comparison with other approaches^10^. If there are differences between the IVW and MR-Egger estimates, this may be due to pleiotropy or heterogeneous treatment effects.

The utility of MR Egger regression hinges upon three core assumptions. First, the INstrument Strength Independent of Direct Effect (InSIDE) assumption requires the effects of SNPs on the exposure and their pleiotropic effects on the outcome to be independent. If the InSIDE assumption holds, estimates for variants with stronger instrument-exposure associations 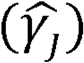 will be closer to the true causal effect parameter than variants with weaker associations^8^. Second, the NO Measurement Error (NOME) assumption requires no measurement error to be present in the instrument-exposure associations, and therefore that the variance of the instrument-exposure association 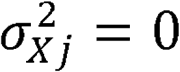. In cases where NOME is strictly satisfied,estimates 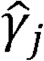 will be equal to γ_j_ and the variance of the ratio estimate for each variant j is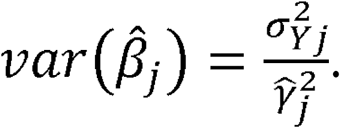

In cases where the NOME assumption is violated, individual variants will suffer from weak instrument bias, leading to attenuation of MR Egger estimates towards the null. This can occur if the SNPs were not genome-wide significant, or were selected from small GWAS. One novel approach to assessing the strength of the NOME assumption is to evaluate the 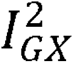statistic, interpreted as the relative degree of attenuation bias in the MR Egger regression in the interval (0,1)^10^. Thus for example, an 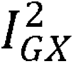 value of 0.7 represents an estimated relative bias of 30% towards the null.

### Weighted median

The weighted median approach is an adaptation of the simple median estimator for two-sample summary MR^13^. For a total number of variants *J*= 2*k* + 1, the simple median approach selects the middle ratio estimate 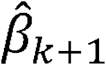, from ordered ratio estimates 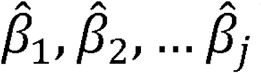^13^. In cases where the total number of variants is even, the median is interpolated as 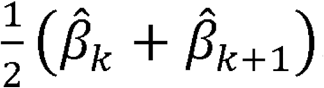. As the simple median approach is inefficient, particularly in cases with variable precision in the set of ratio estimates, it is preferable to incorporate weights in a similar fashion to the IVW and MR Egger approaches. Let 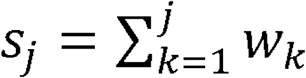 be the sum of weights for the set of variants 1,2, *…j,* standardised so the sum of weights S_j_=1. The weighted median estimator is the median of the distribution of 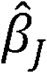 as its 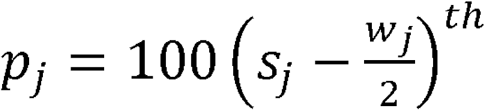 percentile^13^. For the range of percentile values, we perform a linear extrapolation between neighbouring ratio estimates.

An important assumption of the median summary MR approaches is that more than 50% of the genetic variants do not exhibit directional pleiotropy. In the simple median case, this threshold refers to the number of variants, whilst in the weighted median case the 50% threshold is with respect to the weights of the non-pleiotropic variants^13^.

## Implementation

The mrrobust package uses functions from moremata^14^, addplot^15^, and the heterogi^16^ command. For versions of Stata 13 and higher, it can be installed using the.net install command from [http://raw.github.com/remlapmot/mrrobust/master/]. For older versions of Stata, a zip archive of the files is freely available for download at:

[http://github.com/remlapmot/mrrobust].

The package facilitates two-sample summary MR analyses with key features including:

- IVW and MR-Egger regression approaches, including fixed effects MR-Egger regression, standard error correction, and weighting options.
- Unweighted, weighted and penalized weighted median IV estimators, providing pleiotropy robust estimates in cases where fewer than 50% of the genetic instruments are valid.
- Presentation of heterogeneity statistics, and statistics such as 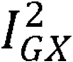 for use in assessing attenuation bias^10^.
- Plotting tools to visualise IVW, MR-Egger and weighted median estimators.
- Illustrative examples and documentation using data from Do et al^17^.

### Applied Examples: Adiposity and Height as predictors of serum glucose levels

To illustrate key features of the mrrobust package, we perform two analyses investigating potential relationships between adiposity, height, and serum glucose. Adiposity was selected owing to the vast body of evidence supporting a positive association with serum glucose levels ^18–21^, whilst height was based upon limited evidence of association ^22–24^. Glucose was selected as an outcome with respect to its hypothesised role in the development of Type-II diabetes^18, 24^

### Applied Example I: Adiposity and Serum Glucose

Though the relationship between adiposity and glucose has received much attention in the literature, such studies are predominantly observational and therefore may be subject to bias from confounding. This provides motivation for considering Mendelian randomization techniques which are able to control for such unobserved confounding. In the initial analysis, we select adiposity as an exposure measured using standardised body mass index (BMI), obtaining estimates of its associations with genotypes and their respective standard errors from Locke et al^25^.

For the outcome, we consider log transformed measures of serum glucose *log(mM)* utilising effect estimates and standard errors from Shin et al^26^. Adopting a GWAS significance p-value threshold of 5x 10^-8^ a total of 79 independent SNPs were identified in both samples. We confirmed the linkage equilibrium (LD) between the SNPs using a clumping algorithm, and a clumping distance of 10000kb, and an LD *R^2^* of 0.001. This resulted in a total of 79 SNPs for use as instrumental variables, details of which are presented in the Web Appendix.

Using mrrobust, we conducted IVW, MR-Egger, and weighted median regression approaches using the above summary data. The code for our analysis is in the Supplementary Material. For IVW and MR Egger approaches the regression was weighted using the variance of the instrument-outcome association. The set of summary MR estimates are presented in Table 1A.

**Table 1.**
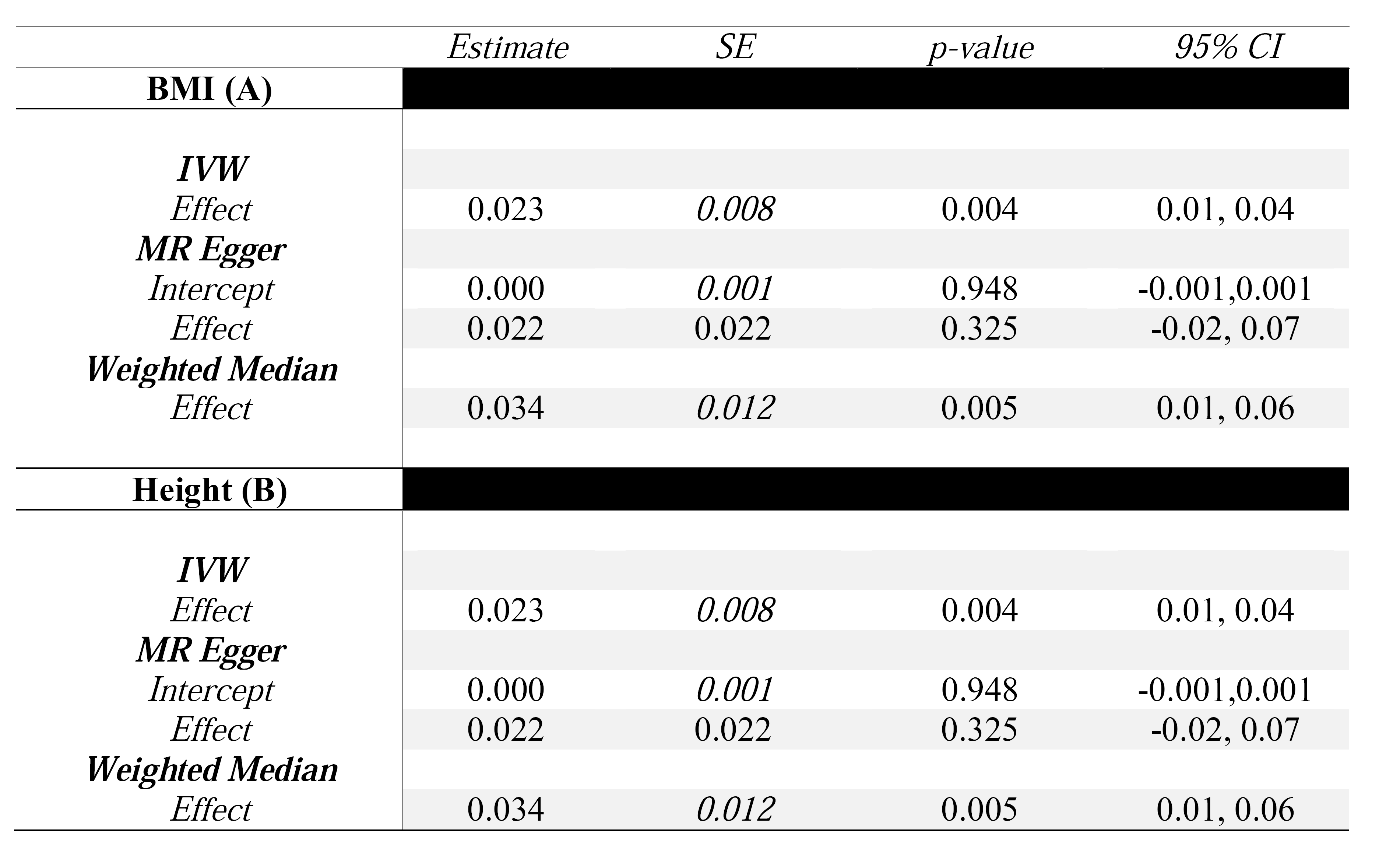
Summary MR estimates for the effect of standardised BMI (A) and height (B) upon log transformed serum glucose

As seen in Table 1A, we find strong evidence of a positive association between BMI and serum glucose using both IVW and weighted median methods. Considering the MR Egger case, a substantial average directional pleiotropic effect was not detected, and the lack of significance with respect to the effect estimate can be attributed to a lack of statistical power. An 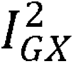 value of 0.88 was reported, which can be interpreted as a relative bias in the MR-Egger estimate of 12% towards the null. The set of estimates from Table 1 are illustrated in Figure 1A using the mreggerplot command.

**Figure 1.**
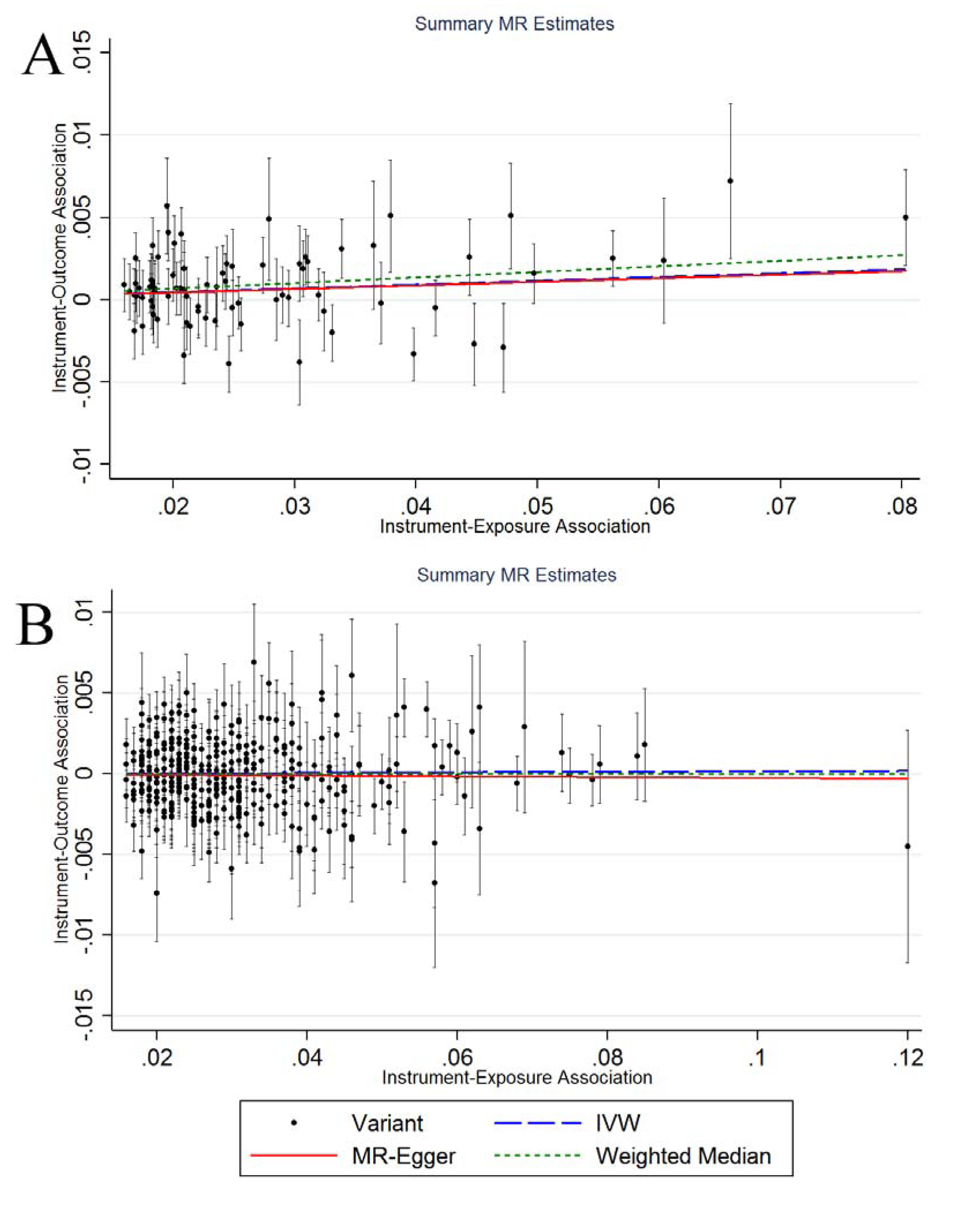
mreggerplot output for applied examples using BMI (A) and Height (B) as exposures.

### Applied Example II: Height and Serum Glucose

As a further example, we consider the effect of standardised height (*meters*) upon serum glucose using summary data from Wood et al^27^, and outcome summary data on log transformed serum glucose from Shin et al^26^. We assess the SNPS for LD using criteria from the previous example, and identify 367 SNPs as suitable instruments for the analysis. The set of summary MR estimates are presented in Table 1B.

From Table 1B we find no evidence against the null hypothesis of no association between height and serum glucose levels using IVW, weighted median, and MR Egger regression. Considering the MR Egger case, there appeared to be no evidence of directional pleiotropy, with an 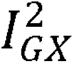 value of 0.90 indicating a relative bias of 10% towards the null. The set of two-sample MR estimates are presented in Figure 1B.

## Discussion

The mrrobust package is a freely available Stata package, containing a number of summary MR estimation methods which can be used to estimate causal effects. In the applied example, the mrrobust package was able to provide a series of estimates, finding evidence of a positive association between BMI and serum glucose, and no evidence of association between height and serum glucose. One possible conclusion that can be drawn from these results is that previously reported associations between height and glucose are driven by confounding factors^28, 29^ It is important, however, to consider the extent to which Mendelian randomization is appropriate for a given analysis, and by extension situations in which mrrobust is suitable.

In the first instance, Mendelian randomization studies only produce unbiased estimates when genetic instruments satisfy the assumptions of each estimator (e.g. IVW, MR-Egger, or weighted median). In two-sample analyses genetic instruments should be associated with the exposure of interest at genome-wide levels of significance (satisfying the first instrumental variable assumption), and pruned for LD to limit the overlap between SNPs. The IVW estimator also requires that genetic variants should not have directional pleiotropic effects. The MR Egger and median estimators are robust to directional pleiotropy if the effects of the exposure are constant. MR Egger regression requires the InSIDE and NOME assumptions. Median methods assume that the number of valid instruments being greater than 50%. In cases where the value of 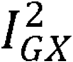 is low, it is possible to use SIMEX or Bayesian error in variables regression methods as methods of correcting for attenuation bias, and these features will be implemented in subsequent releases of the mrrobust package.

In this paper, we have presented the mrrobust Stata package as an accessible toolkit for performing summary MR and instrumental variable analysis using many instruments. It contains a range of summary MR approaches, and should make examining causal relationships using Mendelian randomization more accessible for genetic epidemiologists.

## Supplementary Data

A web appendix containing supplementary materials can be found at:

## Funding

The Medical Research Council (MRC) and the University of Bristol fund the MRC Integrative Epidemiology Unit [MC UU 12013/1, MC UU 12013/9].

## Acknowledgements

The authors would like to thank Michael Holmes, Caroline Dale, Amy Taylor, Rebecca Richmond, Judith Brand, Yanchun Bao, and Kawthar Al-Dabhani for helpful comments on the programs.

## References

1. Davey Smith, G. and S. Ebrahim, ‘Mendelian randomization’: can genetic epidemiology contribute to understanding environmental determinants of disease? International Journal of Epidemiology, 2003. 32(1): p. 1–22.

2. Burgess, S., et al., Mendelian randomization: where are we now and where are we going? Int J Epidemiol, 2015. 44(2): p. 379–88.

3. Evans, D.M. and G. Davey Smith, Mendelian Randomization: New Applications in the Coming Agee of Hypothesis-Free Causality. Annu Rev Genomics Hum Genet, 2015. 16: p. 327–50.

4. Scholder, S.V.K., et al., Mendelian Randomization: The Use of Genes in Instrumental Variable Analyses. Health Economics, 2011. 20(8): p. 893–896.

5. Burgess, S., D.S. Small, and S.G. Thompson, A review of instrumental variable estimators for Mendelian randomization. Stat Methods Med Res, 2015. 0(0): p.1–26.

6. Davies, N.M., et al., The many weak instruments problem and Mendelian randomization. Stat Med, 2015. 34(3): p. 454–68.

7. Burgess, S. and S.G. Thompson, Use of allele scores as instrumental variables for Mendelian randomization. Int J Epidemiol, 2013. 42(4): p. 1134–44.

8. Bowden, J., G. Davey Smith, and S. Burgess, Mendelian randomization with invalid instruments: effect estimation and bias detection through Egger regression. Int J Epidemiol, 2015. 44(2): p. 512–25.

9. Burgess, S., A. Butterworth, and S.G. Thompson, Mendelian randomization analysis with multiple genetic variants using summarized data. Genet Epidemiol, 2013. 37(7): p. 658–65.

10. Bowden, J., et al., Assessing the suitability of summary data for two-sample Mendelian randomization analyses using MR-Egger regression: the role of the I2 statistic. Int J Epidemiol, 2016. 45(6): 1961–1974.

11. Bowden, J., et al., A framework for the investigation of pleiotropy in two-sample summary data Mendelian randomization. Stat Med, 2017. 36(11): p.1783–1802.

12. Yavorska, O.O. and S. Burgess, MendelianRandomization: an R package for performing Mendelian randomization analyses using summarized data. Int J Epidemiol, 2017. https://doi.org/10.1093/ije/dyx034

13. Bowden, J., et al., Consistent Estimation in Mendelian Randomization with Some Invalid Instruments Using a Weighted Median Estimator. Genet Epidemiol, 2016. 40(4): p. 304–14.

14. Jann, B., MOREMATA: Stata module (Mata) to provide various functions. Statistical Software Components, 2005.

15. Jann, B., A note on adding objects to an existing twoway graph. Stata Journal, 2015. 15(3): p. 751–755.

16. Orsini N, H.J., Bottai M, Buchan I., heterogi: Stata module to quantify heterogeneity in a meta-analysis. Statistical Software Components, 2006.

17. Do, R., et al., Common variants associated with plasma triglycerides and risk for coronary artery disease. Nature Genetics, 2013. 45(11): p. 1345-+.

18. Boden, G., Role of fatty acids in the pathogenesis of insulin resistance and NIDDM. Diabetes, 1997. 46(1): p. 3–10.

19. Kim, S.H. and G. Reaven, Obesity and insulin resistance: an ongoing saga. Diabetes, 2010. 59(9): p. 2105–6.

20. O’Gorman, D.J., et al., Exercise training increases insulin-stimulated glucose disposal and GLUT4 (SLC2A4) protein content in patients with type 2 diabetes. Diabetologia, 2006. 49(12): p. 2983–92.

21. Hjerkind, K.V., J.S. Stenehjem, and T.I. Nilsen, Adiposity, physical activity and iisk of diabetes mellitus: prospective data from the population-based HUNT study, Norway. BMJ Open, 2017. 7(1): p. e013142.

22. Ogonowski, J. and T. Miazgowski, Are short women at risk for gestational diabetes mellitus? Eur J Endocrinol, 2010. 162(3): p. 491–7.

23. Brown, D.C., et al., Height and glucose tolerance in adult subjects. Diabetologia, 1991. 34(7): p. 531–3.

24. Asao, K., et al., Short stature and the risk of adiposity, insulin resistance, and type 2 diabetes in middle age: the Third National Health and Nutrition Examination Survey (NHANES III), 1988-1994. Diabetes Care, 2006. 29(7): p. 1632–7.

25. Locke, A.E., et al., Genetic studies of body mass index yield new insights for obesity biology. Nature, 2015. 518(7538): p. 197–U401.

26. Shin, S.Y., et al., An atlas of genetic influences on human blood metabolites. Nat Genet, 2014. 46(6): p. 543–50.

27. Wood, A.R., et al., Defining the role of common variation in the genomic and biological architecture of adult human height. Nat Genet, 2014. 46(11): p. 1173–86.

28. Olatunbosun, S.T. and A.F. Bella, Relationship between height, glucose intolerance, and hypertension in an urban African black adult population: a case for the "thrifty phenotype" hypothesis? J Natl Med Assoc, 2000. 92(6): p. 265–8.

29. Rehunen, S.K.J., et al., Adult height and glucose tolerance: a re-appraisal of the importance of body mass index. Diabet Med, 2017.

